# Benchmarking Protein Language Models for Protein Crystallization

**DOI:** 10.1101/2024.09.02.610758

**Authors:** Raghvendra Mall, Rahul Kaushik, Zachary A. Martinez, Matt W. Thomson, Filippo Castiglione

## Abstract

The problem of protein structure determination is usually solved by X-ray crystallography. Several *in silico* deep learning methods have been developed to overcome the high attrition rate, cost of experiments and extensive trial-and-error settings, for the predicting the crystallization propensities of proteins based on their sequences. In this work, we benchmark the power of open protein language models (PLMs) through the TRILL platform, a bespoke framework democratizing the usage of PLMs for the task of predicting crystallization propensities of proteins.

By comparing LightGBM / XGBoost classifiers built on the embedding representations learned by different PLMs, such as ESM2, Ankh, ProtT5-XL, ProstT5, with the performance of state-of-the-art sequence-based methods like DeepCrystal, ATTCrys and CLPred, we identify the most effective methods for predicting crystallization outcomes. The LightGBM classifiers utilizing embeddings from ESM2 model with 30 and 36 transformer layers and 150 and 3, 000 million parameters respectively have performance gains by 3 *−* 5% then all compared models for various evaluation metrics, including AUPR (Area Under Precision-Recall Curve), AUC (Area Under the Receiver Operating Characteristic Curve), and F1 score on independent test sets.

Furthermore, we fine-tune the ProtGPT2 model available via TRILL to generate crystallizable proteins. Starting with 3, 000 generated proteins and through a step of filtration processes including consensus of all open PLM-based classifiers, sequence identity through CD-HIT, secondary structure compatibility, aggregation screening, homology search and foldability evaluation, we identified a set of 5 novel proteins as potentially crystallizable.

## 1. Introduction

Protein structure at atomic resolution is usually determined by X-ray crystallography [1] or nuclear magnetic resonance (NMR) [2]. However, this is an expensive process where *>* 70% of the total cost is spent on attempts that do not produce crystals of diffraction quality [3]. Crystallization of proteins is a prerequisite for structural determination. Yet, it has been a daunting challenge, with the overall rate of successful attempts ranging between 2 and 10% [4]. The determination of important biological features that help increase the propensity for protein crystallization remains a great challenge. Several machine learning methods and statistical techniques have been developed for sequence-based protein crystallization prediction [5, 6, 7, 8, 9, 10, 11]. These approaches utilize feature-based protein representations including physicochemical and k-mer frequency features from amino acid sequences and corresponding structures. Most of these feature-based techniques undergo a feature selection procedure(s), followed by traditional machine learning techniques such as support vector machines [12, 13], random forests [14] and gradient-boosting machines [15].

The availability of large-scale protein datasets through public databases such as PepcDB [16], enables the use of deep learning techniques for the problem of protein crystallization prediction. DeepCrystal, a deep neural network (DNN) based model was proposed by Elbasir et al. [17] to predict protein crystallization propensity **using just the protein AA sequence as input** without the need to extract additional physio-chemical and k-mer features by implementing convolutional neural networks (CNNs) [18] as backbone. DeepCrystal captures frequently occurring amino acid (AA) k-mers of different lengths driving the crystallization prediction and outperforms state-of-the-art (*sota*) feature-based methods. Furthermore, techniques such as ATTCry [19] design a CNN framework based on multi-scale and multi-head self-attention for crystallization prediction. CLPred [20] uses a bidirectional recurrent neural network with long- and short-term memory (BLSTM) to capture long-range interaction patterns between the k-mers of AA sequence to predict protein crystallizability using the protein AA sequence as input.

A new deep learning pipeline, GCmapCrys [21], was proposed for multistage crystallization propensity prediction by integrating the graph attention networks with the predicted protein contact map. Moreover, it uses BLAST [22] to generate a position-specific scoring matrix, SCRATCH-1D (https://scratch.proteomics.ics.uci.edu/) to use predicted solvent accessibility and secondary structure, and HHblits [23] for multiple sequence alignment (MSA). A similar technique, namely BCrystal [24], utilizes homology, secondary structure, solvent accessibility, torsion angle features in combination with an XGBoost model. However, these techniques, in particular those using MSA are extremely slow (*≈* 30 minutes for one protein sequence) and cannot be used for high-throughput screening of proteins.

Since, the goal of our work was to compare the crystallization propensity of a protein using just their AA sequence and the ability of the model to perform high-throughput screening, hence we focus on methods such as DeepCrystal, ATTCrys and CLPred during our experimental comparisons.

In recent years, application of natural language processing (NLP) methods to protein sequences has led to remarkable breakthroughs for *sota* protein structure and property prediction. The driving force for these breakthroughs is the transformer, a deep learning architecture [25], which uses the concept of self-attention to efficiently capture long-range dependencies and intricate patterns in protein sequences that were previously difficult to discern using traditional deep learning methods [25].

Analogous to using words and sentences to train typical large language models (LLMs), transformer-based models such as ESM2 use individual AAs, peptides, and protein sequences [26] to learn the “language” of life. These protein language models (PLMs) follow a self-supervised learning framework, where the model attempts to predict the identity of randomly masked AAs (usually 15% of the AAs per protein sequence) using the unmasked potions of the protein sequence. For example, ESM2 was pre-trained on the masked language training task with *≈*65 million unique protein sequences from UniRef [26]. After this extensive training, scientists are able to use these pre-trained models to extract high-dimensional representations for their proteins of interest. These vectors can be used for downstream tasks such as protein property prediction, protein clustering, and functional comparisons [27, 28, 29, 24, 30, 31, 32].

In the present work, we perform efficacy assessments of several open source PLMs for the task of predicting protein crystallization using the TRILL platform [33]. TRILL is a comprehensive resource designed to democratize access to *sota* open PLMs, eliminating the requirement for advanced computational skills. Using robust deep learning frameworks such as Pytorch Lightning [34] and HuggingFace Accelerate [35], TRILL provides access to several PLMs such as ESM2 [26], Ankh [36] and ProstT5 [37], specifically for tasks such as protein design and property analysis. Moreover, TRILL facilitates the usage of these PLMs with different model configurations and parameter space. These PLMs in TRILL are complemented by a suite of utilities that enhance user experience and functionality.

In the case of protein sequence classification, the platform provides functionalities to embed protein sequences into vector representations, visualize the embedded protein sequence representation, train custom classifiers, and predict class labels for unseen protein sequences. These diverse tools and functionalities are encapsulated within a command-line interface, organized through ten commands as detailed in the original TRILL paper [33]. In the present work, we utilize the TRILL platform to determine the vector representation of proteins for each PLM using just the AA sequence as input. These vector representations are then passed as training data to classifiers which are optimized through hyper-parameter tuning. This results in optimal crystallization propensity predictor for individual PLM. We then perform a comprehensive comparison of these PLM-based predictors on several independent test sets. Finally, we generate 3000 proteins through a finetuned ProtGPT2 model (on the crystallizable class) and through a series of computational filtration steps identify a reduced set of 5 novel proteins as potentially crystallizable.

The key contributions of the manuscript are:

- Benchmarking different ESM2 models for the task of protein crystallization prediction using raw protein sequences on external balanced, SwissProt and TrEMBL test sets;
- Benchmark PLMs such as Ankh, Ankh-Large, ProstT5 and ProtT5-XL for the task of protein crystallization prediction on external balanced, SwissProt and TrEMBL test sets;
- Comprehensive comparison of open-source PLMs to predict diffraction-quality crystals with superior performance on aforementioned test sets;
- Provide all the code used for benchmarking open-PLMs for crytallization prediction task via github (https://github.com/raghvendra5688/ crystallization_benchmark) for reproducibility and enabling community to utilize TRILL for their protein property prediction task.
- Fine-tune a protein generator namely ProtGPT2 [38] to generate *de novo* protein sequences belonging to the crystallizable class;
- Evaluate, screen and validate the generated proteins to identify a unique set of stable and well-folded proteins.

Figure 1 provides a flow diagram of the proposed framework for predicting protein crystallization propensity.

**Figure 1:**
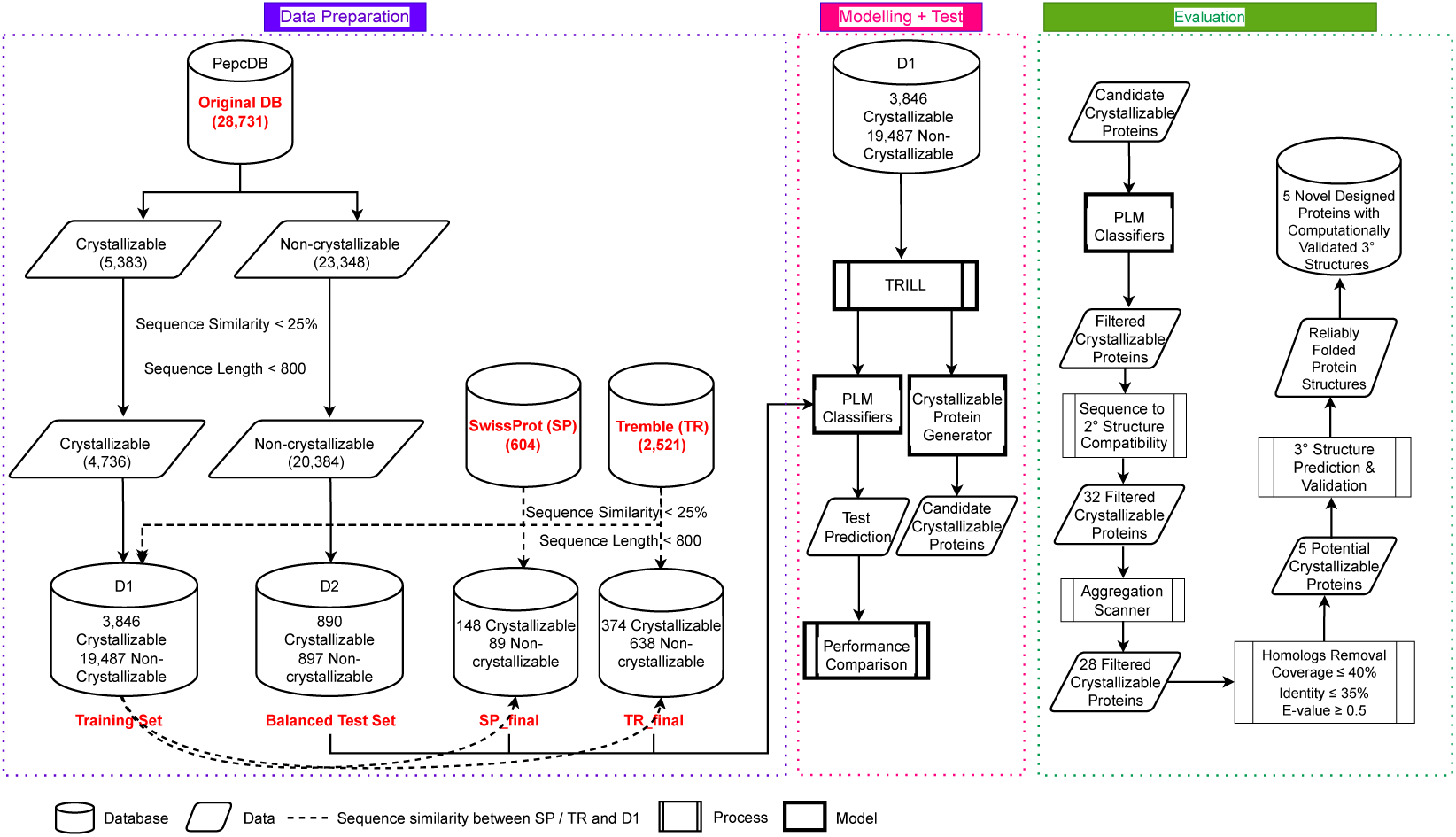
Flowchart of the proposed PLM benchmarking framework for protein crystallization propensity prediction.

## 2. Materials and Methods

### 2.1. Overview

The problem of predicting the crystallization propensity of a protein is a binary classification task. A protein sequence, is given by a sequence of AAs *x* = (*x*_1_*, x*_2_*, …, x_L_*), where *x_i_*, is the *i^th^* amino acid in the sequence and is part of a vocabulary comprising 20 amino acids, while *L* is the length of the protein sequence. A given PLM uses its encoder referred as “tokenizer” (*t*(*·*)) that encodes the AA sequence *x* to an encoded representation (*t*(*x*) *∈* R*^L^*) that is then ingestible for deep learning technique. This is a widely used encoding scheme in natural language processing (NLP) to have a vector representation for words in a sentence [39, 40].

The encoded representation *t*(*x*) is then given as input to the PLM and the final transformer layer of the PLM generates an embedding representation of the protein, preserving meaningful inter-residue relationships and contextual information within the original protein sequence. In mathematical terms *e*(*t*(*x*)) is the embedding of the protein *x*, with *e* : R*^L^ →* R*^d^*, where *d* represents the embedding dimension of the transformer layer of the PLM (note: for comparison reasons, we use different PLMs, thus *d* changes).

Our aim is to learn a function *c*(*·*) that takes as input the embedded protein sequence *e*(*t*(*x*)) and outputs a probability, i.e., *c* : R*^L^ →* [0, 1], where *c*(*·*) is the function computed by the nonlinear classifier. In this work, *c*(*·*) is an XGBoost [41] or a LightGBM [42] classifier.

While fine-tuning individual PLM (either all layers or few layers) with a classification head is an option, some of the PLMs tested in this work are extremely large i.e. ESM2 with 36 transformer layers and *≈* 3 billion parameters. Thus, it is impossible to fine-tune such PLM even with a batch size of 2, given the configuration of the available GPU - NVIDIA RTX A6000 with 48 Gb RAM. Hence, to have a fair evaluation given our GPU capacity, and to understand the learning representation capacity of these PLMs, we considered all these PLMs in a zero-shot learning framework to generate the embedded vector representations for proteins using their AA sequence.

### 2.2. Data Partitioning

We perform our experiment on the processed PepcDB dataset (http://pepcdb.rcsb.org) following the protocols set by Wang et al. [11]. The data set comprises proteins which have been classified into five groups, namely i) diffraction-quality crystals, ii) protein cloning failure, iii) protein material production failure, iv) purification failure, and v) crystallization failure. We consider the proteins labeled as diffraction-quality crystals to be *the* crystallizable class, while other proteins are assigned to the non-crystallizable class. The final dataset comprises 28,731 sequences of which 5,383 proteins belong to the crystallizable class, and the remaining 23,348 are non-crystallizable.

As in [11, 17], all sequences in each class are passed through a filter of sequence identity *>* 25% with other proteins in that class to remove redundant and similar protein sequences within each class.

To divide our dataset into training and test sets, we follow a simple protocol. The maximum length of a protein sequence considered for our model is *L*_max_ = 800. This is done to be compliant with methods like DeepCrystal [17] and CLPred [20], which use the same *L* as the maximum length of the protein sequence. Proteins with *L < L*_max_ are padded with the symbolic representation of gaps. By performing this protein filtering step, the total number of proteins in the dataset is reduced to 25,120.

We follow the procedure used in DeepCrystal [17], ATTCrys [19] and CLPred [20] to divide this dataset into two parts: D_1_ and D_2_ such that D_2_ consists of 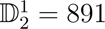 crystallizable and 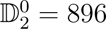 non-crystallizable proteins. Here 1 corresponds to crystallizable and 0 corresponds to non-crystallizable class. Thus, D_2_ represents the fairly balanced test set for performance evaluation as used in DeepCrystal, ATTCrys and CLPred methods. D_1_ has a total of 23,333 protein sequences, where D^1^ = 3, 846 proteins belong to crystallizable class while remaining D^0^ = 19, 487 proteins fall are non-crystallizable.

We also use two independent test sets generated in [1] as external validation sets. The two external datasets, referred as SP final and TR final were obtained from SwissProt and TrEMBL databases respectively, following the protocol detailed in Elbasir et al. [17]. In the SP final dataset, we have 148 proteins belonging to the positive class while remaining 89 sequences are non-crystallizable, whereas in the TR final dataset there are 374 crystallizable proteins and 638 proteins belonging to the negative class. We compare our methods with sota web-servers such as fDETECT [8], DeepCrystal [17], ATTCrys [19] and CLPred [20] on these datasets. For all performance comparison, we provide our test protein sequences to these web-servers to obtain corresponding prediction scores.

### 2.3. Benchmarking Models

The TRILL platform [33] provides access to several PLMs, such as ESM2 [26], Ankh [36], ProstT5 [37] and ProtT5-XL [43], which can generate protein embedding representations via a zero-shot learning framework. Moreover, there are several pretrained PLMs, such as ESM2 [26], ProtGPT2 [38] and ZymCTRL [44], which can either directly generate proteins in a zero-shot fashion or first by fine-tuning these models and then proceed with protein generation. Here we provide a summary of several PLMs used in the present work. For further details of these PLMs, reader’s indulgence is sought.

#### 2.3.1. Evolutionary Scale Modeling (ESM2)

ESM2 is a sota transformer-based protein language model trained on *≈*65 million unique protein sequences [26]. ESM2 has been shown to outperform all tested single-sequence PLMs across a range of structure prediction tasks, enabling atomic resolution structure prediction. While the ESM2 model has been benchmarked for structure prediction, it has not been gauged for protein property prediction and has been shown to not scale for protein function prediction [45]. Moreover, the ESM2 models are available with different architectural configurations, that is, with an increase in number of transformer layers leading to an increase in number of model parameters. The ESM2 models are available with 6, 12, 30, 33 and 36 transformer layers having *≈* 8, 12, 150, 650 and 3,000 million parameters respectively.

#### 2.3.2. Ankh

The Ankh is an optimized general-purpose PLM, as a first version for future specialized high-impact protein modeling tasks. Ankh is pre-trained on the UniRef50 dataset [46], that provides more variability and representation compared to UniRef100 [46] and BFD [47]. The model is tested on a comprehensive set of downstream tasks spanning protein function prediction, structure prediction, and localization prediction. Ankh demonstrated superior performances on the tasks such as fluorescence prediction, solubility prediction, contact prediction, fold prediction, and secondary structure prediction. Additionally, Ankh used Google’s latest TPU v4 hardware and JAX/Flax software for efficient training. Thus, Ankh is presented as a powerful general-purpose PLM that can serve as a foundation for specialized protein modeling tasks, with outstanding performances demonstrated on a wide range of benchmarks. Ankh-Large has *≈*2 billion parameters and is trained using the encoder-decoder architecture, while Ankh base has *<* 10% parameters when compared to the sota models.

##### 2.3.3. ProstT5

ProstT5 is a bilingual language model for protein sequences and structures that leverages the AlphaFold Protein Structure Database (AFDB) [48]. ProstT5 was pre-trained using 34.6 million proteins. It can translate between 1-D amino acid sequences and 1-D structure sequences (3Di tokens). ProstT5 demonstrated the improved performance in various protein function prediction tasks compared to sota sequence-based models such as ProtT5, ESM2 and Ankh. It can perform inverse folding, generate novel AA sequences that adopt a desired structural template, and assess the quality of its own predictions. ProstT5 exemplifies how language modeling techniques and transformers can be used to leverage the wealth of information from protein structure databases such as AFDB. Finally, ProstT5 is a proof-of-concept bilingual PLM that showcases the potential of integrating sequence and structure information for various protein modeling tasks.

##### 2.3.4. ProtT5-XL

ProtT5-XL uses an encoder-decoder framework for training [25]. ProtT5-XL has 3 billion parameters and is trained using an 8-way model parallelism. ProtT5-XL is trained on BFD for 1.2 million steps, followed by a fine-tuning on UniRef50 for 991k steps. Contrary to the original T5 model [48] that masks spans of multiple tokens, ProtT5-XL adopts BERT’s denoising objective to corrupt and reconstruct single tokens using a masking probability of 15%. ProtT5-XL uses the AdaFactor optimizer with inverse square root learning rate schedule for pretraining. Using embeddings from ProtT5-XL as the input to supervised models to predict secondary structure and subcellular localization, it outperformed previous methods on these tasks.

##### 2.3.5. ProtGPT2

ProtGPT2 is a PLM that can generate novel protein sequences which are structurally and functionally similar to natural proteins [38]. ProtGPT2 effectively generates sequences that are distantly related to natural ones but are not a consequence of memorization and repetition. Majority of ProtGPT2 sequences (93%) have significant sequence similarity to natural proteins [38]. AlphaFold predictions show 37% of ProtGPT2 sequences have high confidence (pLDDT *>* 70) for being ordered structures, comparable to 66% for natural sequences. Molecular dynamics simulations indicate ProtGPT2 sequences have similar dynamic properties as natural proteins [38].

Integrating ProtGPT2 sequences into a structural network representation of the protein universe reveals they bridge separate “islands” of known protein structures. ProtGPT2 generates sequences across different structural classes like all-*α*, all-*β*, *α/β*, etc. The model can be conditioned to design proteins for specific families, functions or structural classes. Thus, the unsupervised ProtGPT2 model effectively learns the “protein language” and generates novel sequences that populate unexplored regions of protein structure space while maintaining key structural and functional properties. This highlights the potential of PLMs for *de novo* protein design.

### 2.4. Model Building & Test

We follow a simple protocol to use the TRILL platform for our task of benchmarking PLMs for protein crystallization propensity prediction. Starting with the training sequences *x ∈* D_1_, we obtain embedding representations *e* (*t*(*x*)) for each of the following 9 protein language models: ESM2 T6-8M, ESM2 T12-35M, ESM2 T30-150M, ESM2 T33-650M, ESM2 T36-3B, Ankh, Ankh Large, ProstT5, ProtT5-XL PLMs using the embed function.

The embedding representations *e_k_*(*t*(*x*))*, k* = 1 … 9 are generated in a zero-shot learning setting. These embedding representations of the training set D_1_ are then passed to the XGBoost classifier using the classify utility, where a 10-fold cross-validation technique is used for hyper-parameter optimization. The details of the hyperparameters are available via xgboost classifier script.

The XGBoost classifiers optimizes a weighted average F1-metric during the classification step to address the problem of class-imbalance. We also pass the embedding representations *e_k_*(*t*(*x*)) from each PLM to custom LightGBM models [42] in 10-fold cross-validation setting to generate LightGBM classifiers. We performed a randomized search over a grid of parameters including number of estimators, maximum depth of a tree, number of leaves, minimum child samples, learning rate, subsampling rate, L1 and L2 regularizers during hyper-parameter optimization. The details of the parameter space for LightGBM classifiers are available at hyperparameter tuning script.

Thus, in total we have 9 XGBoost classifiers and 9 LightGBM classifiers, where each classifier is built on top of embedding representations (*e_k_*(*t*(*x*))) obtained from a PLM. After obtaining the XGBoost / LightGBM classifier for each of the 9 PLMs, we pass the test sets to each PLM to obtain embedding representations for the respective set of proteins. Finally, the class label and probability *c* (*e_k_*(*t*(*x*))) for each protein sequence *x* in a given test set and the *k^th^* PLM is obtained by passing its embedding representation *e_k_*(*x*) to the classifier *c*(*·*). We utilize the classify function with ‘–preComputed Embs’ and ‘–preTrained’ utilties in TRILL to obtain the class probability as shown in Figure 2. A consensus of the predictions from the 18 classifiers is obtained by taking average of the probabilities estimated by these classifiers.

**Figure 2:**
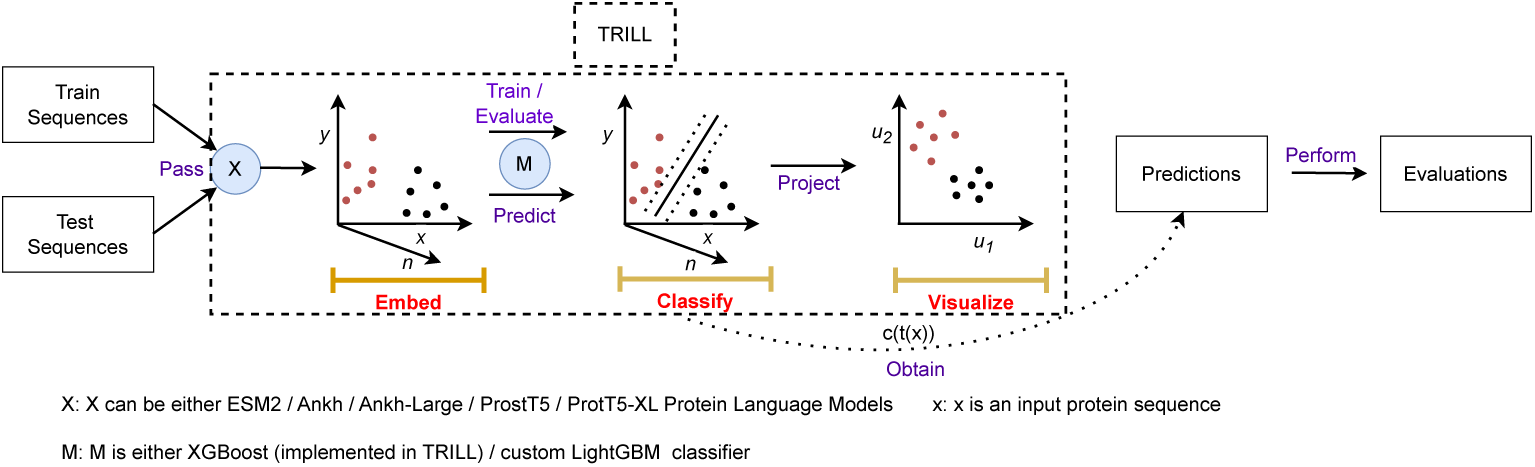
Workflow of building the crystallization propensity prediction classifiers for each PLM and obtaining test set predictions using the TRILL platform. Here the ‘red’ colored dots represent crystallizable proteins and ‘black’ colored dots correspond to non-crystallizable proteins.

A detailed workflow of building the classifiers and obtaining predictions on test sets is highlighted in Figure 2.

### 2.5. Protein Generation

We fine-tune the ProtGPT2 PLM on the crystallizable class (D^1^) using the fine-tune function available in TRILL for 10 epochs. In [33] it was shown that 10 epochs are sufficient to generate synthetic cell penetrating peptides and anti-crispr proteins using ProtGPT2. Thus, the fine-tuned ProtGPT2 model learns the underlying distribution of crystallizable proteins. We then generate a total of 3,000 proteins using the fine-tuned ProtGPT2 model via the lang gen utility. Once we have generated the synthetic proteins, we obtain the embedding representation for the same using the PLMs and visualize these embeddings in a low-dimensional space (2 dimensions) using the visualize function. This function utilizes the Unified Manifold and Approximation (UMAP) algorithm [49] to project the embeddings into a two-dimensional space. Then, the embedding representation for a generated protein is obtained and classified by each of the 18 classifiers. This protein generation and classification process is illustrated in Figure 3. We then follow a series of filtration steps to determine the most promising candidates:

**Figure 3:**
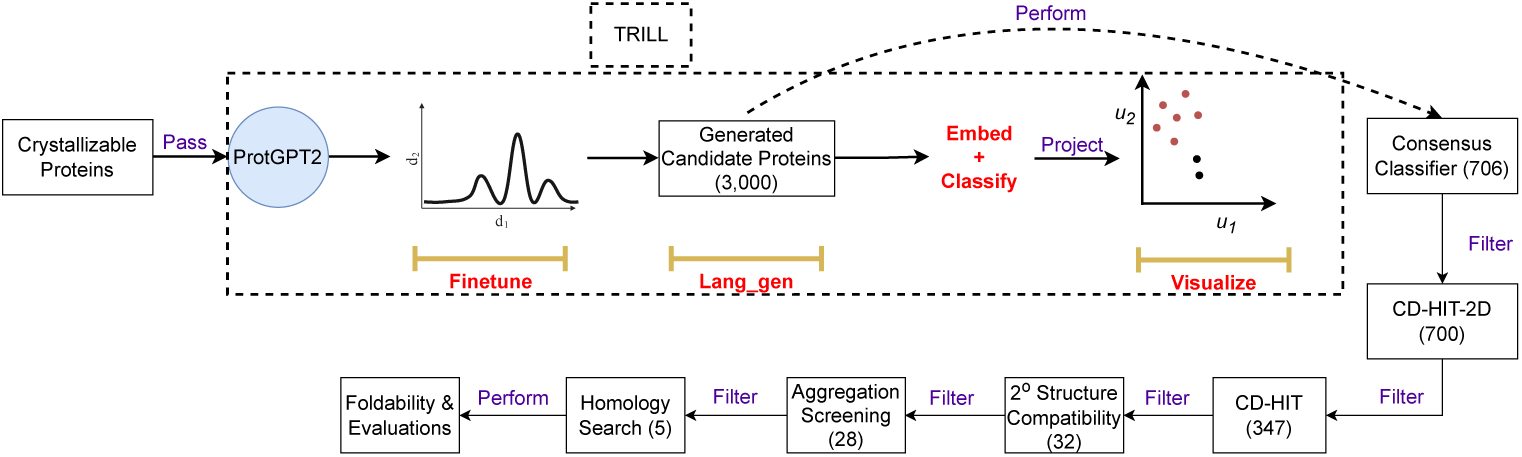
The protocol followed to generate crystallizable proteins using fine-tuned Prot-GPT2 PLM and further downstream filtering and evaluation.

Step 1: A consensus of all PLM-based classifiers consistently identified 706 out of the 3, 000 generated proteins as crystallizable proteins.

Step 2: To remove generated sequences with high sequence identity with training set, we perform CD-HIT-2D [50] with a identity cut-off of *≤* 40%, resulting in 700 protein sequences.

Step 3: CD-HIT is then performed to cluster proteins with *>* 25% sequence identity into groups, leading to a total of 347 proteins with low sequence identity within the group and with the training set.

Step 4: Filtered protein sequences are screened by sequence to secondary structure compatibility scores [51, 52]. The secondary structural characterization of the designed protein sequences is performed by utilizing PSIPRED (standalone ver. 4.02) [53]. This reduces the generated protein set from 347 to 32 candidate sequences.

Step 5: The screened proteins are further evaluated on the basis of presence of aggregation prone regions [54] and 4 sequences are filtered out.

Step 6: The screened proteins are subjected for the availability of any homolog(s) in known protein sequence database, UniRef100 [46], resulting in a reduced set of 5 proteins.

Step 7: The 5 filtered proteins are modeled using a consensus approach by implementing RoseTTAFold2 [55], and AlphaFold2 [56], resulting in 6 model structures (5 from AlphaFold2 and 1 from RoseTTAFold end-2-end prediction) for each protein.

Step 8: Each model structure is refined by implementing GalaxyRefine [57] to generate 5 refined model structures, resulting in 30 candidate model structure for each protein.

Step 9: The modeled structure for each protein are thoroughly analyzed to identify the best model structure (1 out of 30) among the candidate structures using ModFold (ver. 9.0) [23] and ProFitFun [52, 51].

Step 10: Finally, the stereo-chemical quality (all atoms contact and geometry) of the best model structure for each protein is assessed by passing it through ProCheck [58], Errat [59], and MolProbity [60].

By following the aforementioned steps, we filter an initial set of 3, 000 proteins generated from crystallizable class to the set of 5 most likely and high confidence crystallizable proteins.

### 2.6. Evaluation Metrics

The performance of benchmark classifiers is compared with various other sota techniques using quality metrics such as accuracy, Matthew’s correlation coefficient (MCC) as in [31, 17]. We assessed other evaluation metrics, based on TP, TN, false positives (FP) and false negative (FN). We highlight that TP represents the set of proteins which are crystallizable (the true label is 1) and are correctly identified by a given method as crystallizable, i.e., *c* (*e*(*t*(*x*))) *≥* 0.5. Similarly, TN represents the set of proteins which are non-crystallizable (true label is 0) and are correctly identified by a given method as non-crystallizable *c* (*e*(*t*(*x*)))) *<* 0.5. The metrics for evaluation include:

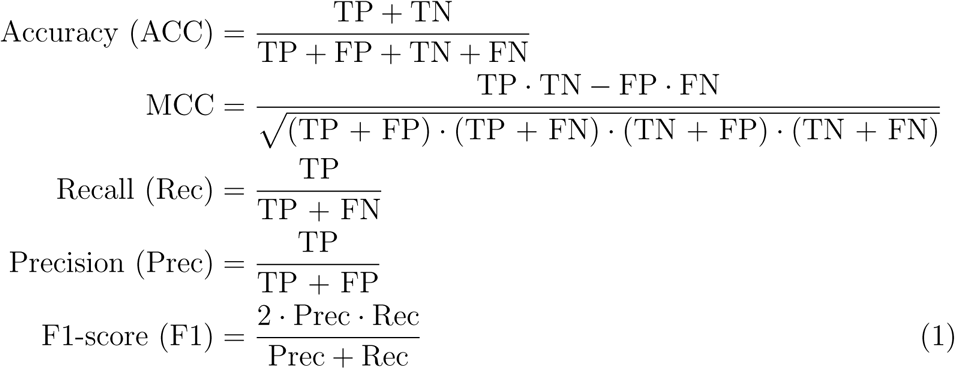

## 3. Experimental Results

We benchmark the predictive performance of the PLMs on the D_2_ test set extracted from the publicly available dataset [11] as described earlier (see Section II *B*). Moreover, we evaluate the quality of predictions from these models on two independent datasets obtained from SwissProt and TrEMBL, the SP final and TR final datasets, respectively. A comprehensive comparison of the PLMs of varying size and configurations including ESM2 T6-8M, ESM2 T12-35M, ESM2 T30-150M, ESM2 T33-650M, ESM2 T36-3B, Ankh, Ankh Large, ProstT5, ProtT5-XL was done against methods like fDETECT, DeepCrystal, ATTCrys and CLPred across these test sets. The evaluation metric values for fDETECT and CLPred were obtained from [17] and [20] respectively. Finally, the cross-validation performance of the XGBoost and LightGBM classifiers built on embedding representations learnt via each PLM on various evaluation metrics is highlighted in Supp. Figures 1 and 2. From Supp. Figures 1 and 2 and Tables 1, 2 and 3, we observe that the XGBoost models are over-fitting on the training set and have poor generalization performance. On the other hand, the LightGBM classifiers have better generalization performance as their cross-validation performance aligns with the performance attained on multiple independent test sets (see Supp. Figure 2 and Tables 1, 2 and 3).

**Table 1:**
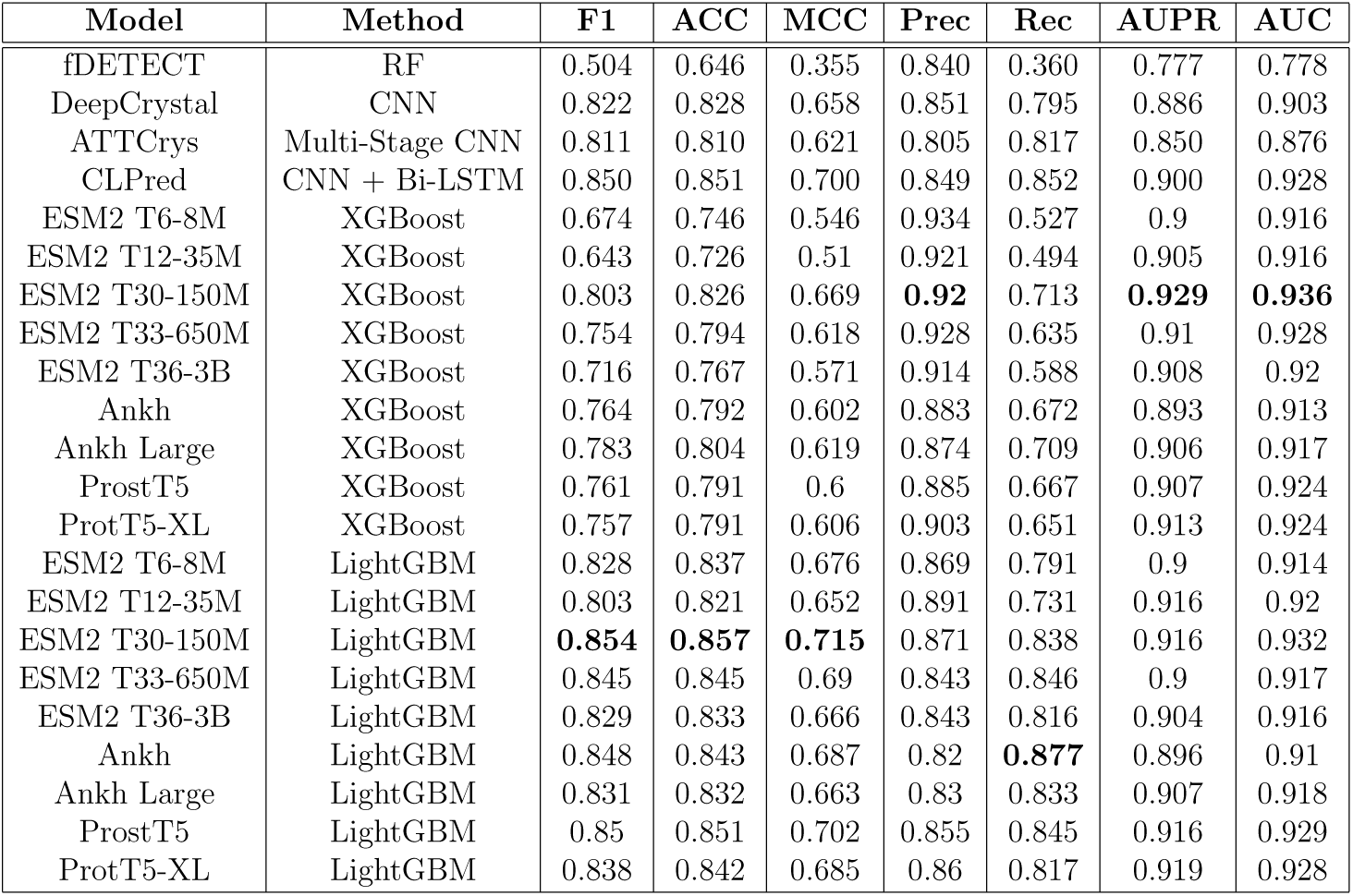
Benchmarking of PLMs in TRILL on the balanced test set against sota methods.

**Table 2:** Benchmarking of PLMs in TRILL on the SP final test set against sota methods.

| Model | Method | F1 | ACC | MCC | Prec | Rec | AUPR | AUC |
| --- | --- | --- | --- | --- | --- | --- | --- | --- |
| fDETECT | RF | 0.580 | 0.616 | 0.381 | 0.913 | 0.425 | 0.882 | 0.837 |
| DeepCrystal | CNN | 0.788 | 0.759 | 0.53 | 0.876 | 0.716 | 0.877 | 0.874 |
| ATTCrys | Multi-Stage CNN | 0.814 | 0.772 | 0.521 | 0.831 | 0.797 | 0.856 | 0.827 |
| CLPred | CNN + Bi-LSTM | 0.832 | 0.801 | 0.599 | 0.885 | 0.783 | 0.880 | 0.887 |
| ESM2 T6-8M | XGBoost | 0.712 | 0.713 | 0.524 | 0.955 | 0.568 | 0.948 | 0.913 |
| ESM2 T12-35M | XGBoost | 0.615 | 0.646 | 0.445 | 0.957 | 0.453 | 0.929 | 0.881 |
| ESM2 T30-150M | XGBoost | 0.836 | 0.814 | 0.646 | 0.933 | 0.757 | 0.947 | 0.919 |
| ESM2 T33-650M | XGBoost | 0.795 | 0.781 | 0.61 | 0.953 | 0.682 | 0.948 | 0.922 |
| ESM2 T36-3B | XGBoost | 0.814 | 0.802 | 0.657 | **0.981** | 0.696 | **0.964** | 0.935 |
| Ankh | XGBoost | 0.761 | 0.743 | 0.528 | 0.907 | 0.655 | 0.932 | 0.906 |
| Ankh Large | XGBoost | 0.84 | 0.819 | 0.653 | 0.934 | 0.764 | 0.955 | 0.93 |
| ProstT5 | XGBoost | 0.829 | 0.81 | 0.648 | 0.948 | 0.736 | 0.957 | **0.94** |
| ProtT5-XL | XGBoost | 0.794 | 0.776 | 0.593 | 0.936 | 0.689 | 0.938 | 0.909 |
| ESM2 T6-8M | LightGBM | 0.871 | 0.848 | 0.694 | 0.924 | 0.824 | 0.953 | 0.915 |
| ESM2 T12-35M | LightGBM | 0.803 | 0.781 | 0.585 | 0.914 | 0.716 | 0.934 | 0.888 |
| ESM2 T30-150M | LightGBM | 0.873 | 0.848 | 0.688 | 0.912 | 0.838 | 0.954 | 0.931 |
| ESM2 T33-650M | LightGBM | 0.883 | 0.857 | 0.699 | 0.901 | 0.865 | 0.946 | 0.921 |
| ESM2 T36-3B | LightGBM | **0.911** | **0.89** | **0.769** | 0.924 | **0.899** | 0.961 | 0.938 |
| Ankh | LightGBM | 0.885 | 0.857 | 0.694 | 0.885 | 0.885 | 0.931 | 0.912 |
| Ankh Large | LightGBM | 0.876 | 0.848 | 0.681 | 0.894 | 0.858 | 0.954 | 0.929 |
| ProstT5 | LightGBM | 0.898 | 0.878 | 0.751 | 0.941 | 0.858 | **0.964** | **0.94** |
| ProtT5-XL | LightGBM | 0.873 | 0.848 | 0.688 | 0.912 | 0.838 | 0.952 | 0.927 |

**Table 3:** Benchmarking of PLMs in TRILL on the TR final test set against sota methods.

| Model | Method | F1 | ACC | MCC | Prec | Rec | AUPR | AUC |
| --- | --- | --- | --- | --- | --- | --- | --- | --- |
| fDETECT | RF | 0.747 | 0.841 | 0.663 | 0.918 | 0.631 | 0.768 | 0.887 |
| DeepCrystal | CNN | 0.781 | 0.841 | 0.657 | 0.800 | 0.762 | 0.815 | 0.910 |
| ATTCrys | Multi-Stage CNN | 0.758 | 0.810 | 0.605 | 0.718 | 0.802 | 0.793 | 0.880 |
| CLPred | CNN + Bi-LSTM | 0.807 | 0.854 | 0.690 | 0.787 | 0.829 | 0.865 | 0.930 |
| ESM2 T6-8M | XGBoost | 0.729 | 0.835 | 0.648 | 0.926 | 0.602 | 0.911 | 0.944 |
| ESM2 T12-35M | XGBoost | 0.692 | 0.819 | 0.616 | 0.932 | 0.551 | 0.901 | 0.939 |
| ESM2 T30-150M | XGBoost | 0.816 | 0.875 | 0.73 | 0.9 | 0.746 | **0.933** | **0.96** |
| ESM2 T33-650M | XGBoost | 0.772 | 0.854 | 0.685 | 0.912 | 0.668 | 0.917 | 0.954 |
| ESM2 T36-3B | XGBoost | 0.783 | 0.863 | 0.708 | **0.94** | 0.671 | 0.925 | 0.955 |
| Ankh | XGBoost | 0.756 | 0.839 | 0.649 | 0.858 | 0.676 | 0.875 | 0.932 |
| Ankh Large | XGBoost | 0.797 | 0.858 | 0.69 | 0.844 | 0.754 | 0.898 | 0.942 |
| ProstT5 | XGBoost | 0.762 | 0.84 | 0.65 | 0.846 | 0.693 | 0.88 | 0.943 |
| ProtT5-XL | XGBoost | 0.776 | 0.852 | 0.678 | 0.878 | 0.695 | 0.91 | 0.948 |
| ESM2 T6-8M | LightGBM | 0.846 | 0.885 | 0.755 | 0.841 | 0.85 | 0.909 | 0.947 |
| ESM2 T12-35M | LightGBM | 0.807 | 0.868 | 0.712 | 0.873 | 0.751 | 0.9 | 0.941 |
| ESM2 T30-150M | LightGBM | **0.862** | **0.894** | **0.778** | 0.833 | 0.893 | 0.929 | 0.959 |
| ESM2 T33-650M | LightGBM | 0.829 | 0.867 | 0.723 | 0.787 | 0.877 | 0.901 | 0.944 |
| ESM2 T36-3B | LightGBM | **0.862** | **0.894** | 0.777 | 0.835 | 0.89 | 0.925 | 0.956 |
| Ankh | LightGBM | 0.82 | 0.853 | 0.706 | 0.748 | **0.906** | 0.869 | 0.928 |
| Ankh-Large | LightGBM | 0.835 | 0.87 | 0.732 | 0.785 | 0.89 | 0.892 | 0.944 |
| ProstT5 | LightGBM | 0.839 | 0.875 | 0.739 | 0.799 | 0.882 | 0.903 | 0.949 |
| ProtT5-XL | LightGBM | 0.844 | 0.88 | 0.749 | 0.814 | 0.877 | 0.912 | 0.951 |

### 3.1. Balanced Test Set Results

On the balanced test set consisting of 1787 proteins (891 crystallizable and 896 non-crystallizable), the ESM2 T30-150M PLM (with LightGBM classifier) achieves a prediction accuracy of 85.7%. This is better than the current sota method, CLPred (85.1%). The ESM2 T30-150M (LightGBM) also reaches the best performance of 0.854 and 0.715 for quality metrics such as F1 score and MCC, respectively, as observed from Table 1. These quality metrics take into account the class imbalance in the data set. The performance of ESM2 T30-150M (LightGBM) is 0.4% and 1.5% better in absolute terms than the current sota sequence-based crystallization predictor i.e., CLPred. Moreover, ESM2 T30-150M is 3.2%, 2.9%, and 5.7% better than DeepCrystal for F1 score, accuracy, and MCC metrics, respectively.

However, with respect to quality metrics such as AUPR and AUC, the ESM2 T30-150M (with XGBoost classifier) model leads when compared to all other benchmark models as observed from Table 1 and Figures 4a, 4b, 5a, and 5b. The ESM2 T30-150M (XGBoost) model reaches AUPR = 0.929 and AUC = 0.936. This is 4.3% and 3.3% better than DeepCrystal for AUPR and AUC metrics, respectively, as observed in Table 1. Furthermore, from Table 1, we observe that PLMs with XGBoost classifier available via TRILL tend to handle the class-imbalance worse than PLMs with custom LightGBM classifier. This is highlighted from the superior performance of PLMs with LightGBM classifier on F1-score and MCC metrics when compared to their equivalent XGBoost classifiers available via TRILL as depicted in Table 1. Overall, PLMs trained with either LightGBM classifier outperform CLPred, ATTCrys and DeepCrystal across all metrics on balanced test set.

**Figure 4:**
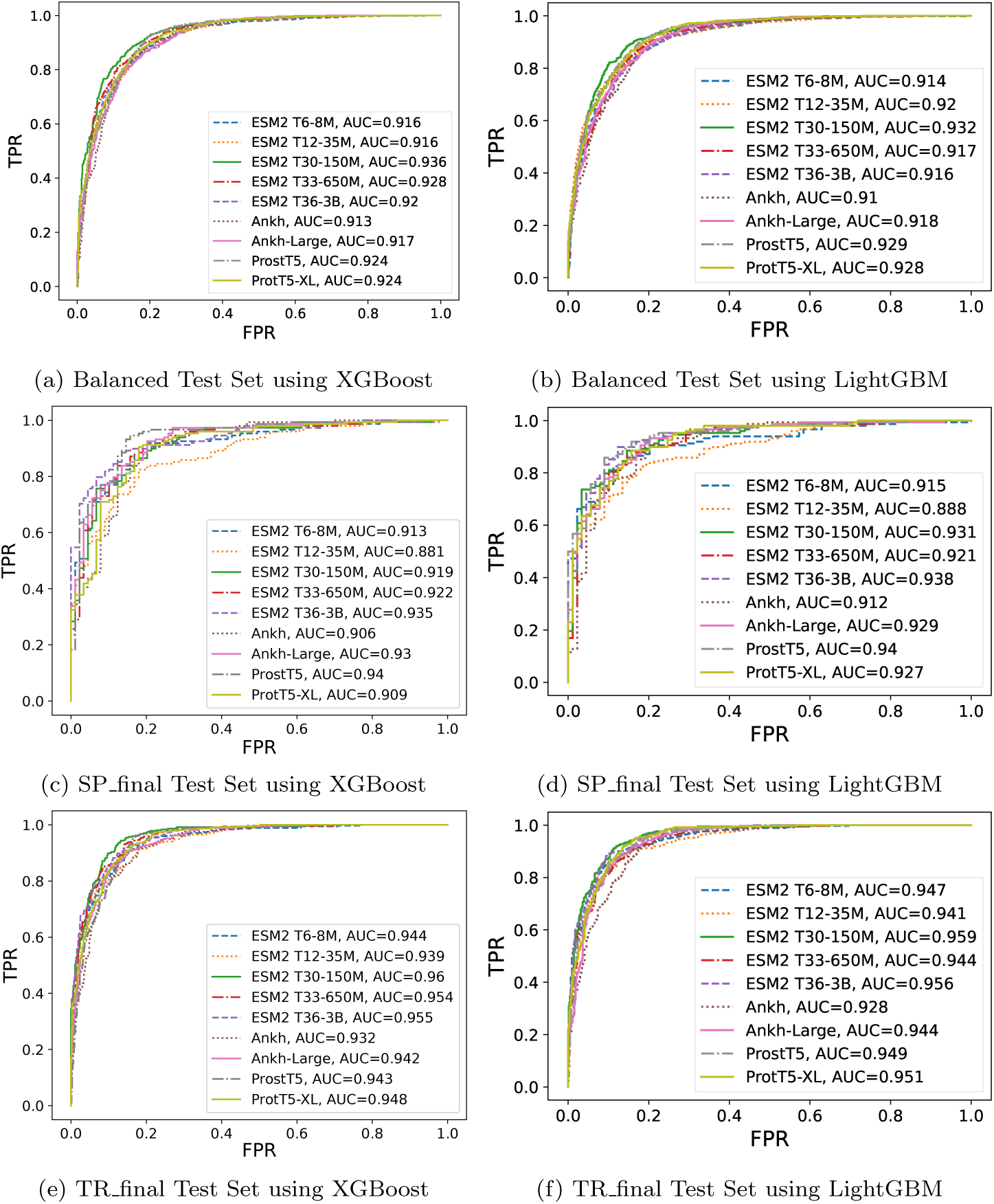
Comparison of area under receiver operating curve (AUC) of benchmark PLMs for the crystallization prediction task across the three different test sets. (a) AUC for fairly balanced test set using XGBoost, (b) AUC for SP_final dataset using XGBoost, (c) AUC for TR final dataset using XGBoost, (d) AUC for fairly balanced test set using LightGBM, (e) AUC for SP_final dataset using LightGBM, and (f) AUC for TR_final dataset using LightGBM.

**Figure 5:**
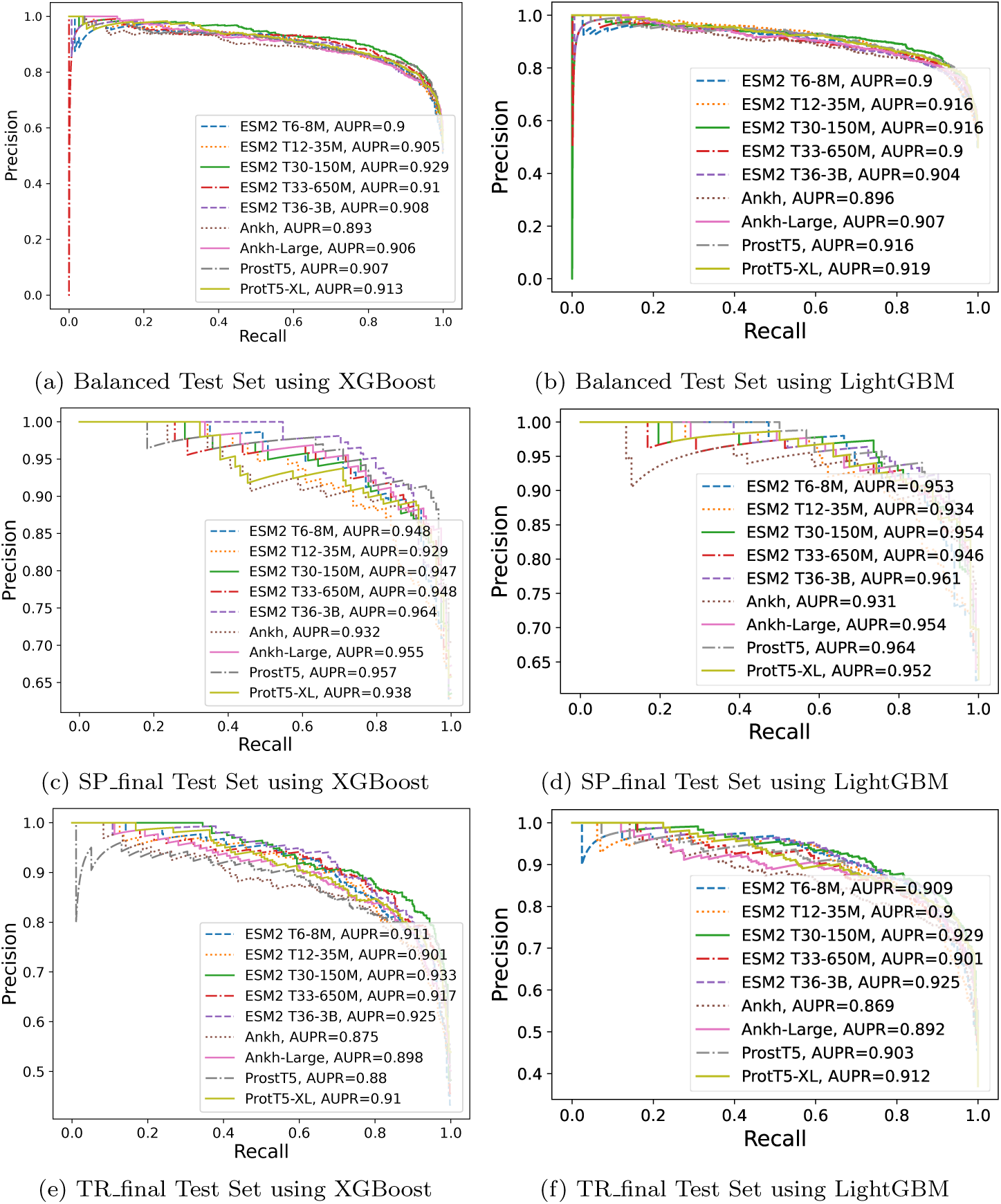
Comparison of area under precision-recall curve (AUPR) of benchmark PLMs for the crystallization prediction task across the three different test sets. (a) AUPR for fairly balanced test set using XGBoost, (b) AUPR for SP_final dataset using XGBoost, (c) AUPR for TR_final dataset using XGBoost, (d) AUPR for fairly balanced test set using LightGBM, (e) AUPR for SP_final dataset using LightGBM, and (f) AUPR for TR_final dataset using LightGBM.

### 3.2. SP final Test Set Results

A second experiment is performed on the reduced SP final dataset obtained from SP Pre dataset [1]. The ESM2 T36-3B model (with LightGBM classifier) outperforms sota sequence-based crystallization predictors like CL-Pred and DeepCrystal for the majority of the metrics, including F1, accuracy, MCC and precision as depicted in Table 2. The ESM2 T36-3B (LightGBM) model also outperforms other PLMs available via TRILL for these quality metrics as shown in Table 2. ESM2 T36-3B model (LightGBM) achieves a prediction accuracy of 89%, which is 9% and 14% better than CLPred and DeepCrystal respectively (see Table 2). From Table 1, we observe ESM2 T36-3B model (LightGBM) attains an MCC of 0.769 and F1-score of 0.911, whereas CLPred obtains an MCC of 0.599 and F1-score of 0.832 indicating 17% and 8% improvement in performance. The ProstT5 model (with Light-GBM classifier) achieves the best AUC (0.940) and AUPR (0.964) compared to other PLM-based classifiers as depicted in Figures 4c, 4d, 5c and 5d.

We observe from Table 2 that small sized ESM2 models such as ESM2 T6-12M and ESM2 T12-35M cannot outperform CLPred for several quality metrics but bigger sized ESM2 models easily surpass sota models like fDETECT, DeepCrystal, ATTCrys and CLPred. Finally, the SP final test set comprises 237 proteins with very little sequence similarity with training set and still ESM2 T36-3B classifiers (desgined with XGBoost / LightGBM) out-performs majority of sequence-based predictors on several evaluation metrics highlighting their effectiveness for crystallization propensity prediction.

### 3.3. TR final Test Set Results

We perform a final experiment to test for crystallization propensities of proteins using sota crystallization tools and benchmark PLM-based classifiers available via TRILL platform on the TR final dataset [1]. ESM2 T30-150M model (with LightGBM classifier) achieves a prediction accuracy of 89.4%, which is 4% better than CLPred (85.4%), 5.3% better than DeepCrystal (84.1%) and fDETECT (84.1%). It is also 0.9% better than the next-best ESM2 T6-8M (LightGBM) model that attains an accuracy of 88.5% as depicted in Table 3. The ESM2 T30-150M model (LightGBM) achieves the best F1 (0.862) and MCC (0.778) as highlighted in Table 3 and second best performance for AUC (0.929) and AUPR (0.959) when compared to ESM2 T30-150M (XGBoost), which achieves an AUC of 0.933 and AUPR of 0.960 as indicated in Table 3 and Figures 4e, 4f, 5e and 5f.

Interestingly, we observe from Table 3 that LightGBM classifiers are superior than their counterpart XGBoost classifiers for the same PLM models and configurations highlighting their generalization capability (see Supp. Figure 2). Finally, on the TR final dataset comprising 1012 proteins (far more than SP final test set), the PLM-based classifiers are superior than DeepCrystal, ATTCrys and CLPred w.r.t. several evaluation metrics.

### 3.4. Protein Generation Results

The selected crystallizable candidates (*n* = 347) were trimmed on the basis of sequence to secondary structural compatibility (CS-Score *≥* 40 and CSS-Scores *≥* 20) [52, 51], resulting in a dataset of 32 proteins. The cut-off values for CS- and CSS-Scores were adopted from their benchmarking of successfully designed proteins [51]. These proteins were further tapered to 28 proteins, based on presence of aggregation protein region screening [61], and to 5 proteins based on screening against UniRef100 [46].

The proteins with pairwise sequence coverage *≥* 40%, sequence identity *≥* 35% and e-value *≤* 0.5 were discarded while screening for available homolog(s) in known protein sequence database (UniRef100), resulting in the set of 5 proteins. These protein were modeled by implementing RoseTTAFold (end-2-end prediction; 1 candidate structure for each protein) [62] and AlphaFold2 (*n* = 5 candidate structures for each protein) [56], followed by structure refinement by using GalaxyRefine (*n* = 30; 5 refined candidate structures for each candidate structure) [63]. The best model structure for each protein, selected on the basis of consensus score from ModFold [23] and ProFitFun [52]. The best model structure for each protein along with the distribution of backbone di-hedrals (Ramachandran Map) are depicted in Figure 6. A summary of different quality assessment statistics of the best model structures is provided in Table 4. Additionally, the predicted Global Distance Test -Template Score (GDT-TS), Template Modeling Score (TMS), Global Quality Score (GQS), and Average Quality Score (OAQS) for all the candidate model structures are provided in Table 4.

**Figure 6:**
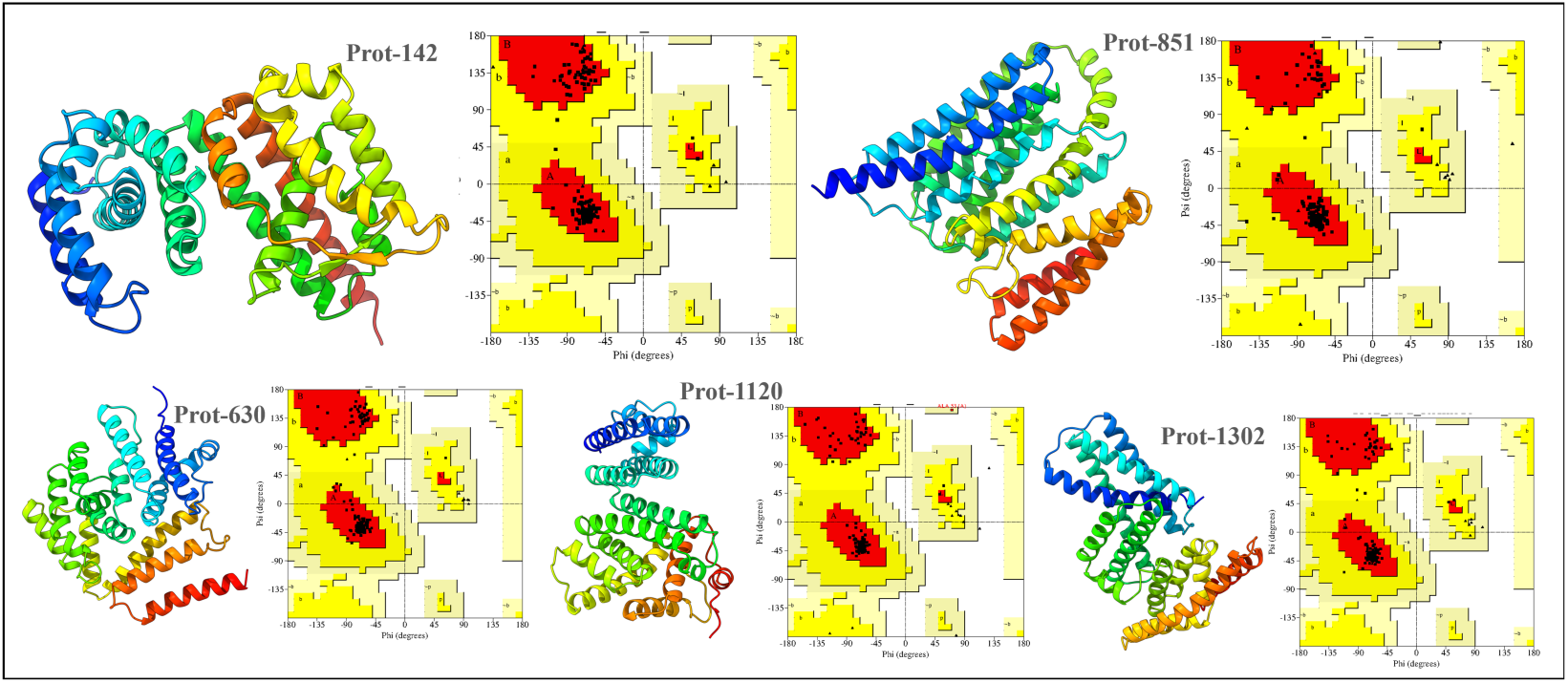
Best model structures for the 5 candidate proteins identified through our crystallizable protein generator workflow.

**Table 4:** Summary of different quality evaluation parameters for the best model structure for each of the selected protein.

| Quality Parameters | Prot-142 | Prot-630 | Prot-851 | Prot-1120 | Prot-1302 |
| --- | --- | --- | --- | --- | --- |
| Ramachandran Distribution |  |  |  |  |  |
| Core Region | 98.9% | 98.1% | 98.8% | 98.5% | 97.4% |
| Allowed Region | 1.1% | 1.9% | 1.2% | 1.1% | 2.6% |
| Generously Allowed Region | 0.0% | 0.0% | 0.0% | 0.4% | 0.0% |
| Disallowed Region | 0.0% | 0.0% | 0.0% | 0.0% | 0.0% |
| Bond Lengths within Limits | 96.6% | 97.3% | 96.6% | 97.9% | 96.5% |
| Bond Angles within Limits | 93.8% | 94.3% | 92.6% | 94.0% | 92.9% |
| Planner Groups within Limits | 98.8% | 100.0% | 98.1% | 100.0% | 99.0% |
| Favored Rotamers | 98.7% | 98.6% | 97.8% | 97.1% | 99.5% |
| Errat Score | 97.3% | 99.0% | 98.1% | 98.0% | 96.4% |
| MolProbity Score | 1.72 | 1.21 | 1.57 | 1.31 | 1.40 |
| Predicted GDT-TS Score | 0.75 | 0.84 | 0.63 | 0.72 | 0.88 |
| Predicted TM-Score | 0.83 | 0.83 | 0.65 | 0.80 | 0.82 |
| Global Quality Score | 0.44 | 0.41 | 0.45 | 0.44 | 0.35 |
| Average Quality Score | 0.67 | 0.69 | 0.58 | 0.65 | 0.69 |

The quality metrics for the best model structure of selected proteins (Prot-142, Prot-630, Prot-851, Prot-1120, and Prot-1302) ensured the accuracy of the tertiary structure prediction (Table 4). For all the model structures, the Ramachandran distribution of backbone di-hedral angles (*ϕ* and *ψ*) is found to be distributed in the allowed regions, mainly the core region (colored ‘red’), as shown in Table 4 and Figure 6. The predicted model structure for Prot-630 and Prot-1302 had the highest quality score (=0.69), followed by Prot-142 (=0.67), Prot-1120 (=0.65), and Prot-851 (=0.58). Notably, the predicted GDT-TS (0.84 for Prot-630 and 0.88 for Prot-1302) and predicted TM Score (0.83 for Prot-630 and 0.82 for Prot-1302) for these protein structure fall in the highly reliable range for predicted model structure (0.8 - 1.0). The GDT-TS and TM Score varies from 0-1, where 1 shows the highest level of structural prediction. The relative predicted quality of the model structure for Prot-851 was observed to be lower as compared to the model structures of other proteins. The secondary and tertiary structures of the selected protein revealed them to be mainly *α*-proteins, except for Prot-142 that has some fraction of residues (about 4%) resulting into *β*-strands.

The functional annotations including biological processes (BP), molecular functions (MF) and cellular components (CC) associated with the generated proteins are provided in Supp. Table 1. Additionally, the two proteins with the maximum functional annotations were Prot-1120 and Prot-1302. The functional annotations associated with these proteins is depicted in Figure 7. We observed that Prot-142 and Prot-630 are localized in cytoplasm, associated to different membranes such as cellular anatomical entity and mainly involved in different metabolic processes and bio-synthetic processes as depicted in Supp. Table 1. The designed protein, Prot-851, while being associated with plasma membrane and cell peripheries such as cellular anatomical entity, was predicted to perform diverse transporter activities by its involvement in different metabolic and transport processes. In contrast to the functional characterization of Prot-142, Prot-630, and Prot-851, the designed proteins Prot-1120 and Prot-1302 were predicted to be involved in the highly diverse set of molecular functions and biological processes as illustrated in Figure 7. For instance, Prot-1120, with the similar cellular localization of other designed proteins, was predicted to be involved in a wider range of metabolic processes, viz. phosphorous, phosphate-containing, and organo-nitrogen compound metabolic processes, primary and cellular metabolic processes, and overall regulation of cellular processes. The Prot-1120 was predicted to be involved catalytic activity, calcium-dependent phospholipid binding, transferase activity, purine ribonucleoside triphosphate binding, small molecule binding, phosphoric ester hydrolase activity, ion binding, organic cyclic compound binding, carbohydrate derivative binding, and heterocyclic compound binding. Further, Prot-1302 is computationally characterized to perform metabolic and biosynthetic process along with trans-membrane transport of various compounds. With the involvement in a diverse set of biological processes, the Prot-1302 was predicted to perform ion channel activity, ATP binding, trans-membrane transporter activity, transferase activity, phosphotransferase activity, purine ribonucleoside triphosphate binding, small molecules and ions binding, organic cyclic compound binding, carbohydrate derivative binding, and heterocyclic compound binding.

**Figure 7:**
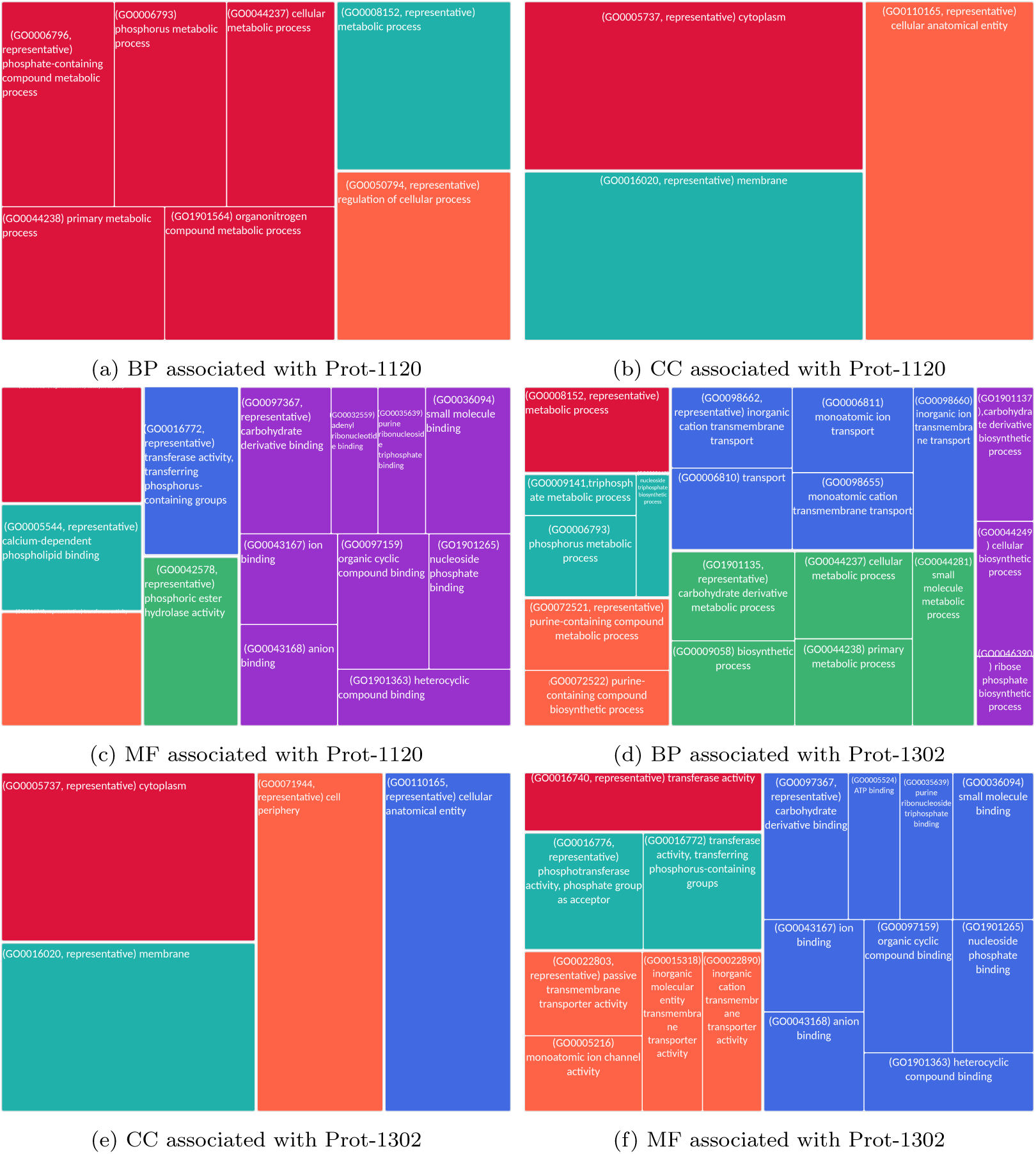
Functional annotations associated with Prot-1120 and Prot-1302.

With a comprehensive computational functional characterization, we believe that experimental validation of Prot-1120 and Prot-1302 can lead to the novel functional proteins that can be fine-tuned to have desired functions.

## 4. Discussion & Conclusion

One of the main challenges for protein structure determination is that only about 2-10% of pursued protein targets yield high-resolution protein structures [64]. Upon investigating these estimates in the TargetDB database [6], it was observed that among the 150, 727 cloned targets that were deposited into TargetDB, only 37, 398 (24.8%) were successfully purified, 12, 923 (8.6%) further successfully crystallized, and 6,942 (4.6%) resulted in diffraction quality crystals [65]. Additionally, majority of the cost of structure determination is consumed by the failed attempts [7] as crystallization is a process that is characterized by a significant rate of attrition. The reasons for this attrition include the need for the crystals to be sufficiently large (*>* 50 micrometers), pure in composition, regular in structure, and without significant internal imperfections. Furthermore, to produce diffraction-quality crystals, an empirical or trial-and-error approach is commonly used, in which a large number of experiments are brute-forced to find a suitable setup [66], often resulting in failure. Thus, the above provides strong motivation to develop accurate and efficient *in silico* sequence-based protein crystallization predictors that allow high-throughput screening of candidate protein sequences for favorable crystallization propensity.

In this paper, we benchmark open-PLMs accessed via the TRILL platform, a framework enabling democratization of protein language models, for sequence-based protein crystallization propensity prediction. The main objective is to determine whether PLMs trained on hundreds of millions of protein sequences can discriminate crystallizable proteins from non-crystallizable ones without fine-tuing using just the raw protein sequences as input. These PLMs encode the raw protein sequences and generate embedding (vector) representations. We then built optimized tree-based classifiers (XGBoost / LightGBM) on top of these embedding representations to estimate their discriminative capacity without the need to manually engineered biological and physiochemical features. By implementing a thorough benchmarking on a set of independent test sets, we observe that these open-PLM based classifiers consistently outperform state-of-the-art deep learning techniques, such as DeepCrystal, ATTCrys and CLPred, on several evaluation metrics.

DeepCrystal [17] captures frequent amino acid *k*-mers in the input sequence using a set of parallel convolution filters of varying sizes with the CNN design providing the freedom of calculating local dependencies with different filter sizes. Conversely, CLPred [20] uses a BiLSTM deep learning architecture to capture high-order, long-range interaction patterns between *k*-mers making it better than the CNN-based DeepCrystal as indicated in Tables 1, 2 and 3. However, open source protein language models trained on several million protein sequences are much better than smaller and crystallization specific deep learning models like DeepCrystal, ATTCrys and CLPred (see Tables 1, 2 and 3), even with no additional fine-tuning and a simple linear probing approach i.e. building classifiers on top of embedding representations. In particular, the **ESM2 T30-150M** and **ESM2 T36-3B** based models (with LightGBM classifier) outperform every other benchmark model on the three independent test sets for quality metrics such as *F1-score*, *accuracy*, *MCC*, and *precision*.

This success can be attributed to the huge amount of data on which these PLMs are trained, the underlying transformer architecture which can capture local and long-range contextual dependencies in protein sequences through attention mechanism [25] and generate meaningful and discriminative embedding representations for the downstream crystallization task.

The proposed methodology illustrates its ability to generate and filter unique crystallizable proteins as well as engineer proteins to achieve desired properties and functions. These proteins may aid in the better understanding of biological processes, as well as the rapid development of new medicines and materials. For example, a designed protein with certain mutations could aid in understanding the roles of specific amino acid residue(s) in the natural protein. Similarly, protein-based therapeutic regimes that involve improvements in the efficacy, stability, solubility, or specificity of certain enzymes, antibodies, and hormones may be accelerated with computational engineering with the help of proposed workflow. Furthermore, computational design may help in the development of more efficient, stable, and selective enzymes that can considerably boost industrial output in the fields of bio-catalysis, food industry, and bio-fuels.

## 6. Acknowledgements

The authors would like to acknowledge Dr. Thomas Launey for his valuable feedback which helped to better position the paper.

## 7. Author Contributions

R.M., M.T. and F.C. conceived the study. R.M. and R.K. performed the data curation. R.M., Z.M. and R.K. designed the methodology. R.M. and R.K. performed the experiments and visualizations. All authors contributed in writing, reviewing and editing the manuscript.

## References

[1] H. Wang, J. Wang, How cryo-electron microscopy and x-ray crystallography complement each other, Protein Science 26 (2017). URL https://api.semanticscholar.org/CorpusID:5167928

[2] K. Wüthrich, Protein structure determination in solution by nmr spectroscopy., Journal of Biological Chemistry 265 (36) (1990) 22059–22062.

[3] R. Service, Structural genomics, round 2, Science 307 (5715) (2005) 1554–1558. arXiv:https://www.science.org/doi/pdf/10.1126/science.307.5715.1554, doi:10.1126/science.307.5715.1554. URL https://www.science.org/doi/abs/10.1126/science.307.5715.1554

[4] T. C. Terwilliger, D. I. Stuart, S. Yokoyama, Lessons from structural genomics., Annual review of biophysics 38 (2009) 371–83. URL https://api.semanticscholar.org/CorpusID:23040480

[5] J. Gao, Z. Wu, G. Hu, K. Wang, J. Song, A. J. Joachimiak, L. Kurgan, Survey of predictors of propensity for protein production and crystallization with application to predict resolution of crystal structures., Current protein & peptide science 19 2 (2017) 200–210. URL https://api.semanticscholar.org/CorpusID:19536343

[6] J. Hu, K. Han, Y. Li, J. yu Yang, H. Shen, D.-J. Yu, Targetcrys: protein crystallization prediction by fusing multi-view features with two-layered svm, Amino Acids 48 (2016) 2533–2547. URL https://api.semanticscholar.org/CorpusID:3709016

[7] L. Kurgan, A. Razib, S. Aghakhani, S. Dick, M. J. Mizianty, S. Jahandideh, Crystalp2: sequence-based protein crystallization propensity prediction, BMC Structural Biology 9 (2009) 50 – 50. URL https://api.semanticscholar.org/CorpusID:10007933

[8] F. Meng, C. Wang, L. Kurgan, fdetect webserver: fast predictor of propensity for protein production, purification, and crystallization, BMC Bioinformatics 18 (2017). URL https://api.semanticscholar.org/CorpusID:23971370

[9] M. J. Mizianty, L. Kurgan, Cryspred: accurate sequence-based protein crystallization propensity prediction using sequence-derived structural characteristics., Protein and peptide letters 19 1 (2012) 40–9. URL https://api.semanticscholar.org/CorpusID:6091681

[10] S. Jahandideh, A. Mahdavi, Rfcrys: sequence-based protein crystallization propensity prediction by means of random forest., Journal of theoretical biology 306 (2012) 115–9. URL https://api.semanticscholar.org/CorpusID:795568

[11] H. Wang, M. Wang, H. Tan, Y. Li, Z. Zhang, J. Song, Predppcrys: Accurate prediction of sequence cloning, protein production, purification and crystallization propensity from protein sequences using multi-step heterogeneous feature fusion and selection, PLoS ONE 9 (2014). URL https://api.semanticscholar.org/CorpusID:1013246

[12] H. Drucker, C. J. C. Burges, L. Kaufman, A. Smola, V. N. Vapnik, Support vector regression machines, in: Neural Information Processing Systems, 1996. URL https://api.semanticscholar.org/CorpusID:743542

[13] J. A. K. Suykens, T. V. Gestel, J. D. Brabanter, B. D. Moor, J. Vandewalle, Least squares support vector machines, 2002. URL https://api.semanticscholar.org/CorpusID:53940663

[14] L. Breiman, Random forests, Machine Learning 45 (2001) 5–32. URL https://api.semanticscholar.org/CorpusID:89141

[15] J. H. Friedman, Greedy function approximation: A gradient boosting machine., Annals of Statistics 29 (2001) 1189–1232. URL https://api.semanticscholar.org/CorpusID:39450643

[16] A. Kouranov, L. Xie, J. de la Cruz, L. Chen, J. Westbrook, P. E. Bourne, H. M. Berman, The rcsb pdb information portal for structural genomics, Nucleic acids research 34 (suppl 1) (2006) D302–D305.

[17] A. Elbasir, B. Moovarkumudalvan, K. Kunji, P. R. Kolatkar, R. Mall, H. Bensmail, Deepcrystal: a deep learning framework for sequence-based protein crystallization prediction, Bioinformatics 35 (13) (2019) 2216–2225.

[18] Y. LeCun, L. Bottou, Y. Bengio, P. Haffner, Gradient-based learning applied to document recognition, Proceedings of the IEEE 86 (11) (1998) 2278–2324.

[19] C. Jin, J. Gao, Z. Shi, H. Zhang, Attcry: Attention-based neural network model for protein crystallization prediction, Neurocomputing 463 (2021) 265–274.

[20] W. Xuan, N. Liu, N. Huang, Y. Li, J. Wang, Clpred: a sequence-based protein crystallization predictor using blstm neural network, Bioinformatics 36 (Supplement 2) (2020) i709–i717.

[21] P.-H. Wang, Y.-H. Zhu, X. Yang, D.-J. Yu, Gcmapcrys: integrating graph attention network with predicted contact map for multi-stage protein crystallization propensity prediction, Analytical Biochemistry 663 (2023) 115020.

[22] S. F. Altschul, T. L. Madden, A. A. Schäffer, J. Zhang, Z. Zhang, W. Miller, D. J. Lipman, Gapped blast and psi-blast: a new generation of protein database search programs, Nucleic acids research 25 (17) (1997) 3389–3402.

[23] L. J. McGuffin, S. M. A. Alharbi, Modfold9: a web server for independent estimates of 3d protein model quality, Journal of Molecular Biology (2024). URL https://api.semanticscholar.org/CorpusID:268433093

[24] A. Elbasir, R. Mall, K. Kunji, R. Rawi, Z. Islam, G.-Y. Chuang, P. R. Kolatkar, H. Bensmail, Bcrystal: an interpretable sequence-based protein crystallization predictor, Bioinformatics 36 (5) (2020) 1429–1438.

[25] A. Vaswani, N. Shazeer, N. Parmar, J. Uszkoreit, L. Jones, A. N. Gomez, Ł. Kaiser, I. Polosukhin, Attention is all you need, Advances in neural information processing systems 30 (2017).

[26] Z. Lin, H. Akin, R. Rao, B. Hie, Z. Zhu, W. Lu, N. Smetanin, R. Verkuil, O. Kabeli, Y. Shmueli, et al., Evolutionary-scale prediction of atomic-level protein structure with a language model, Science 379 (6637) (2023) 1123–1130.

[27] M. Bernhofer, B. Rost, Tmbed: transmembrane proteins predicted through language model embeddings, BMC bioinformatics 23 (1) (2022) 326.

[28] E. Goffinet, R. Mall, A. Singh, R. Kaushik, F. Castiglione, Mate-pred: Multimodal attention-based tcr-epitope interaction predictor, bioRxiv (2024) 2024– 01.

[29] R. Mall, A. Elbasir, H. Almeer, Z. Islam, P. R. Kolatkar, S. Chawla, E. Ullah, A modeling framework for embedding-based predictions for compound–viral protein activity, Bioinformatics 37 (17) (2021) 2544–2555.

[30] R. Mall, Solxplain: An explainable sequence-based protein solubility predictor, BioRxiv (2019) 651067.

[31] R. Rawi, R. Mall, K. Kunji, C.-H. Shen, P. D. Kwong, G.-Y. Chuang, Parsnip: sequence-based protein solubility prediction using gradient boosting machine, Bioinformatics 34 (7) (2018) 1092–1098.

[32] S. Khurana, R. Rawi, K. Kunji, G.-Y. Chuang, H. Bensmail, R. Mall, Deepsol: a deep learning framework for sequence-based protein solubility prediction, Bioinformatics 34 (15) (2018) 2605–2613.

[33] Z. A. Martinez, R. M. Murray, M. W. Thomson, Trill: Orchestrating modular deep-learning workflows for democratized, scalable protein analysis and engineering, bioRxiv (2023).

[34] W. Falcon, J. Borovec, A. Wälchli, N. Eggert, J. Schock, J. Jordan, N. Skafte, V. Bereznyuk, E. Harris, T. Murrell, et al., Pytorchlightning/pytorchlightning: 0.7. 6 release, Zenodo (2020).

[35] S. Gugger, L. Debut, T. Wolf, P. Schmid, Z. Mueller, S. Mangrulkar, M. Sun, B. Bossan, Accelerate: Training and inference at scale made simple, efficient and adaptable., https://github.com/huggingface/accelerate (2022).

[36] A. Elnaggar, H. Essam, W. Salah-Eldin, W. Moustafa, M. Elkerdawy, C. Rochereau, B. Rost, Ankh: Optimized protein language model unlocks general-purpose modelling, arXiv preprint arXiv:2301.06568 (2023).

[37] M. Heinzinger, K. Weissenow, J. G. Sanchez, A. Henkel, M. Steinegger, B. Rost, Prostt5: Bilingual language model for protein sequence and structure, bioRxiv (2023) 2023–07.

[38] N. Ferruz, S. Schmidt, B. Höcker, Protgpt2 is a deep unsupervised language model for protein design, Nature communications 13 (1) (2022) 4348.

[39] Q.-s. Zhang, S.-C. Zhu, Visual interpretability for deep learning: a survey, Frontiers of Information Technology & Electronic Engineering 19 (1) (2018) 27–39.

[40] X. Zhang, J. Zhao, Y. LeCun, Character-level convolutional networks for text classification, Advances in neural information processing systems 28 (2015).

[41] T. Chen, C. Guestrin, Xgboost: A scalable tree boosting system, in: Proceedings of the 22nd acm sigkdd international conference on knowledge discovery and data mining, 2016, pp. 785–794.

[42] G. Ke, Q. Meng, T. Finley, T. Wang, W. Chen, W. Ma, Q. Ye, T.-Y. Liu, Lightgbm: A highly efficient gradient boosting decision tree, Advances in neural information processing systems 30 (2017).

[43] A. Elnaggar, M. Heinzinger, C. Dallago, G. Rehawi, Y. Wang, L. Jones, T. Gibbs, T. Feher, C. Angerer, M. Steinegger, et al., Prottrans: Toward understanding the language of life through self-supervised learning, IEEE transactions on pattern analysis and machine intelligence 44 (10) (2021) 7112– 7127.

[44] G. Munsamy, S. Lindner, P. Lorenz, N. Ferruz, Zymctrl: a conditional language model for the controllable generation of artificial enzymes, in: NeurIPS Machine Learning in Structural Biology Workshop, 2022.

[45] F.-Z. Li, A. P. Amini, Y. Yue, K. K. Yang, A. X. Lu, Feature reuse and scaling: Understanding transfer learning with protein language models, bioRxiv (2024) 2024–02.

[46] B. E. Suzek, Y. Wang, H. Huang, P. B. McGarvey, C. H. Wu, U. Consortium, Uniref clusters: a comprehensive and scalable alternative for improving sequence similarity searches, Bioinformatics 31 (6) (2015) 926–932.

[47] A. Elnaggar, M. Heinzinger, C. Dallago, G. Rihawi, Y. Wang, L. Jones, T. Gibbs, T. Feher, C. Angerer, M. Steinegger, et al., Prottrans: Towards cracking the language of life’s code through self-supervised deep learning and high performance computing. arxiv 2020, arXiv preprint arXiv:2007.06225 (2007).

[48] K. Tunyasuvunakool, J. Adler, Z. Wu, T. Green, M. Zielinski, A. Žídek, A. Bridgland, A. Cowie, C. Meyer, A. Laydon, et al., Highly accurate protein structure prediction for the human proteome, Nature 596 (7873) (2021) 590–596.

[49] L. McInnes, J. Healy, J. Melville, Umap: Uniform manifold approximation and projection for dimension reduction, arXiv preprint arXiv:1802.03426 (2018).

[50] W. Li, A. Godzik, Cd-hit: a fast program for clustering and comparing large sets of protein or nucleotide sequences, Bioinformatics 22 (13) (2006) 1658–1659.

[51] R. Kaushik, K. Y. J. Zhang, A protein sequence fitness function for identifying natural and nonnatural proteins, Proteins: Structure 88 (2020) 1271 – 1284. URL https://api.semanticscholar.org/CorpusID:218659828

[52] R. Kaushik, K. Y. J. Zhang, Profitfun: a protein tertiary structure fitness function for quantifying the accuracies of model structures, Bioinformatics (2021). URL https://api.semanticscholar.org/CorpusID:237572876

[53] D. W. A. Buchan, D. T. Jones, The psipred protein analysis workbench: 20 years on, Nucleic Acids Research 47 (2019) W402 – W407. URL https://api.semanticscholar.org/CorpusID:149920922

[54] R. Kaushik, T. Launey, Decoding protein aggregation through computational approach: Identification and scoring of aggregation-prone regions in protein sequences, bioRxiv (2024). arXiv:https://www.biorxiv.org/content/early/2024/06/12/2024.06.11.598423.full.pdf, doi:10.1101/2024.06.11.598423. URL https://www.biorxiv.org/content/early/2024/06/12/2024.06. 11.598423

[55] B. M, D. F, A. I, D. J, O. S, L. Gr, W. J, C. Q, K. Ln, S. Rd, M. C, P. H, A. C, G. Cr, D. A, P. Jh, R. Av, A. van Dijk, E. Ac, O. Dj, S. T, B. C, P.-K. T, R. Mk, D. U, Y. Ck, B. Je, G. Kc, G. Nv, A. Pd, R. Rj, B. D, Accurate prediction of protein structures and interactions using a 3-track neural network, Science (New York, N.Y.) 373 (2021) 871 – 876. URL https://api.semanticscholar.org/CorpusID:236141270

[56] J. M. Jumper, R. Evans, A. Pritzel, T. Green, M. Figurnov, O. Ronneberger, K. Tunyasuvunakool, R. Bates, A. Žídek, A. Potapenko, A. Bridgland, C. Meyer, S. A. A. Kohl, A. Ballard, A. Cowie, B. Romera-Paredes, S. Nikolov, R. Jain, J. Adler, T. Back, S. Petersen, D. Reiman, E. Clancy, M. Zielinski, M. Steinegger, M. Pacholska, T. Berghammer, S. Bodenstein, D. Silver, O. Vinyals, A. W. Senior, K. Kavukcuoglu, P. Kohli, D. Hassabis, Highly accurate protein structure prediction with alphafold, Nature 596 (2021) 583 – 589. URL https://api.semanticscholar.org/CorpusID:235959867

[57] G. R. Lee, J. Won, L. Heo, C. Seok, Galaxyrefine2: simultaneous refinement of inaccurate local regions and overall protein structure, Nucleic acids research 47 (W1) (2019) W451–W455.

[58] R. Laskowski, M. MacArthur, J. Thornton, Procheck: validation of protein-structure coordinates. international tables for crystallography, Vol F Chapter 21 (4) (2012) 684–687.

[59] C. Colovos, T. O. Yeates, Verification of protein structures: patterns of non-bonded atomic interactions, Protein science 2 (9) (1993) 1511–1519.

[60] R. Kaushik, K. Y. Zhang, An integrated protein structure fitness scoring approach for identifying native-like model structures, Computational and Structural Biotechnology Journal 20 (2022) 6467–6472.

[61] V. Cima, A. Kunka, E. Grakova, J. Planas-Iglesias, M. Havlasek, M. Subramanian, M. Beloch, M. Marek, K. Slaninova, J. Damborsky, Z. Prokop, D. Bednar, J. Martinovic, Prediction of aggregation prone regions in proteins using deep neural networks and their suppression by computational design, bioRxiv (2024). arXiv:https://www.biorxiv.org/content/early/2024/ 03/11/2024.03.06.583680.full.pdf, doi:10.1101/2024.03.06.583680. URL https://www.biorxiv.org/content/early/2024/03/11/2024.03. 06.583680

[62] R. Krishna, J. Wang, W. Ahern, P. Sturmfels, P. Venkatesh, I. Kalvet, G. R. Lee, F. S. Morey-Burrows, I. V. Anishchenko, I. R. Humphreys, R. McHugh, D. Vafeados, X. Li, G. A. Sutherland, A. Hitchcock, C. N. Hunter, A. Kang, E. Brackenbrough, A. K. Bera, M. Baek, F. DiMaio, D. Baker, Generalized biomolecular modeling and design with rosettafold all-atom, bioRxiv (2023). URL https://api.semanticscholar.org/CorpusID:264039660

[63] L. Heo, H. Park, C. Seok, Galaxyrefine: protein structure refinement driven by side-chain repacking, Nucleic Acids Research 41 (2013) W384–W388. URL https://api.semanticscholar.org/CorpusID:16307376

[64] R. F. Service, Structural genomics, round 2, Science 307 (2005) 1554 – 1558. URL https://api.semanticscholar.org/CorpusID:29088350

[65] L. Kurgan, M. J. Mizianty, Sequence-based protein crystallization propensity prediction for structural genomics: Review and comparative analysis, Natural Science 1 (2009) 93–106. URL https://api.semanticscholar.org/CorpusID:18564654

[66] N. E. Chayen, Turning protein crystallisation from an art into a science., Current opinion in structural biology 14 5 (2004) 577–83. URL https://api.semanticscholar.org/CorpusID:26208535

