## Supplementary for "Benchmarking Protein Language Models for Protein Crystallization"

### Benchmarking Protein Language Models for Protein Crystallization using TRILL

Mall et al

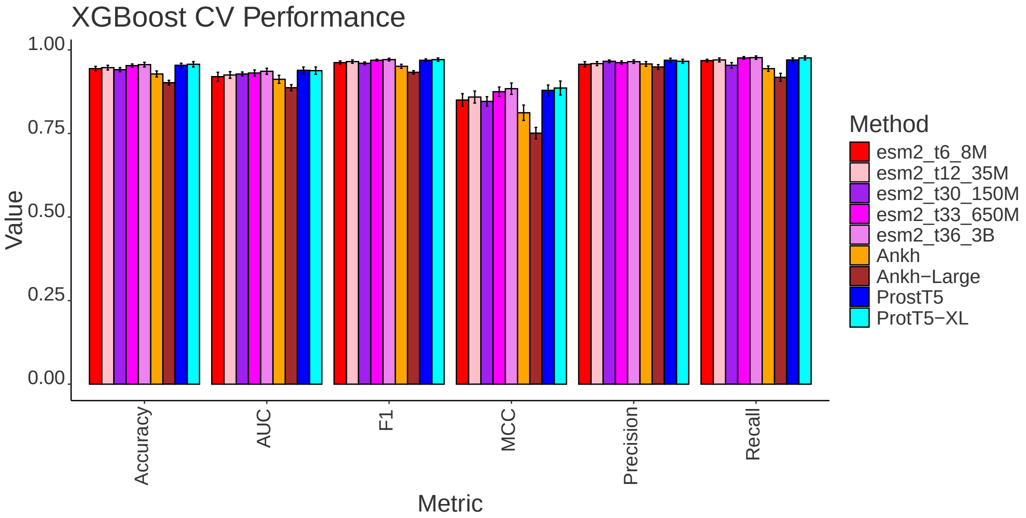


Figure 1: Cross-validation performance of XGBoost classifiers built from different PLMs.

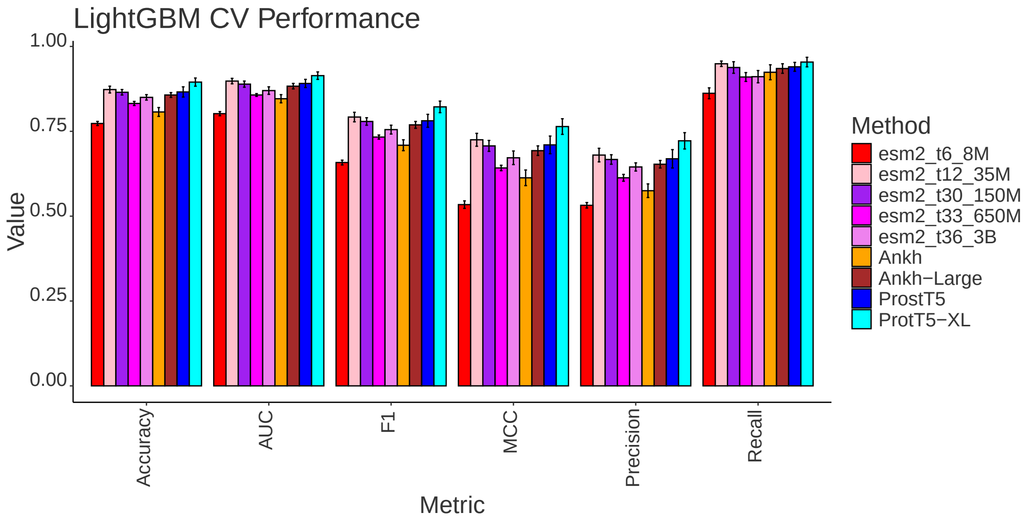


Figure 2: Cross-validation performance of LightGBM classifiers built from different PLMs.

| Protein | Cellular Component | Biological Process | Molecular Functions |
|---|---|---|---|
| Prot-142 | GO:0005737, GO:0016020, GO:0110165 | GO:0006810, GO:0008152, GO:0009058, GO:0044237, GO:0044238, GO:0044249, GO:0044281, GO:0050794, GO:1901564 | |
| Prot-630 | GO:0005737, GO:0110165 | GO:0008152, GO:0009987, GO:0044237, GO:0044238 | GO:0003824 |
| Prot-851 | GO:0005886, GO:0016020, GO:0071944, GO:0110165 | GO:0006810, GO:0009987, GO:0044237 | GO:0008324, GO:0015291, GO:0015318, GO:0022804, GO:0022890 |
| Prot-1120 | GO:0005737, GO:0016020, GO:0110165 | GO:0006793, GO:0006796, GO:0008152, GO:0044237, GO:0044238, GO:0050794, GO:1901564 | GO:0003824, GO:0005544, GO:0016740, GO:0016772, GO:0032559, GO:0035639, GO:0036094, GO:0042578, GO:0043167, GO:0043168, GO:0097159, GO:0097367, GO:1901265, GO:1901363 |
| Prot-1302 | GO:0005737, GO:0016020, GO:0071944, GO:0110165 | GO:0006793, GO:0006810, GO:0006811, GO:0008152, GO:0009058, GO:0009141, GO:0009142, GO:0044237, GO:0044238, GO:0044249, GO:0044281, GO:0046390, GO:0072521, GO:0072522, GO:0098655, GO:0098660, GO:0098662, GO:1901135, GO:1901137 | GO:0005216, GO:0005524, GO:0015318, GO:0016740, GO:0016772, GO:0016776, GO:0022803, GO:0022890, GO:0035639, GO:0036094, GO:0043167, GO:0043168, GO:0097159, GO:0097367, GO:1901265, GO:1901363 |

Table 1: Sequence and structure based functional annotation of selected designed protein.
